# Production of biopolymer precursors beta-alanine and L-lactic acid from CO_2_ with metabolically versatile *Rhodococcus opacus* DSM 43205

**DOI:** 10.1101/2020.08.26.267856

**Authors:** Laura Salusjärvi, Leo Ojala, Gopal Peddinti, Michael Lienemann, Paula Jouhten, Juha-Pekka Pitkänen, Mervi Toivari

## Abstract

Hydrogen oxidizing autotrophic bacteria are promising hosts for CO_2_ conversion into chemicals. In this work, we engineered the metabolically versatile lithoautotrophic bacterium *Rhodococcus opacus* strain DSM 43205 for synthesis of polymer precursors. Aspartate decarboxylase (panD) or lactate dehydrogenase (ldh) were expressed for beta-alanine or L-lactic acid production, respectively. The heterotrophic cultivations on glucose produced 25 mg L^-1^ beta-alanine and 742 mg L^-1^ L-lactic acid, while autotrophic cultivations with CO_2_, H_2_ and O_2_ resulted in the production of 1.8 mg L^-1^ beta-alanine and 146 mg L^-1^ L-lactic acid. Beta-alanine was also produced at 345 µg L^-1^ from CO_2_ in electrobioreactors, where H_2_ and O_2_ were provided by water electrolysis. This work demonstrates that *R. opacus* DSM 43205 can be readily engineered to produce chemicals from CO_2_ and provides base for its further metabolic engineering.

## Introduction

Tackling global warming requires shifting to carbon-neutral chemical manufacture. A disruptive technology for carbon-neutral chemical production is the third generation (3G) biorefining using microbial hosts for CO_2_ conversion into chemicals with renewable energy (Liu et al. 2020). Genetic engineering of autotrophic microbes allows assembling synthetic production routes for direct conversion of CO_2_ into valuable compounds (Hu et al. 2019). Particularly attractive for 3G biorefining is microbial conversion of CO_2_ into valuable compounds in dark conditions using energy from H_2_ oxidation, as H_2_ can be provided through water electrolysis using renewable energy source (e.g. solar panels). Two main CO_2_-fixing, H_2_-oxidizing alternatives are strictly anaerobic acetogenic bacteria such as *Clostridium ljungdahlii* and aerobic lithoautotrophic bacteria such as *Cupriavidus necator*. The anaerobic acetogenic bacteria use the Wood-Ljungdahl pathway for CO_2_ fixation. This pathway requires less energy to fix CO_2_ than the Calvin-Benson-Bassham cycle used by aerobic lithoautotrophic bacteria (Emerson and Stephanopoulos 2019). However, the acetogens are limited to ATP supply with the Wood-Ljungdahl pathway, which may lead to depletion of intracellular acetyl-CoA and restriction of both growth and synthesis of energy-intense products (Molitor et al. 2017). In contrast, the aerobic lithoautotrophic bacteria can couple the oxidation of H_2_ to the electron transfer chain with O_2_ as the final electron acceptor for the respirative ATP generation. Therefore, the aerobic lithoautotrophic bacteria can generate more energy to produce biomass and complex natural products such as poly-hydroxyalkanoates (PHA) (Teixeira et al. 2018; Brigham 2019).

The gram-negative *C. necator* H16 is the most studied aerobic facultatively lithoautotrophic bacterium, and has already been engineered to produce chemicals from CO_2_ (Yu 2018; Brigham 2019). Typically, mg L^-1^ level product titers for e.g. isopropanol, 2-hydroxyisobutyric acid, methyl ketones, alkanes, α-humulene and isobutanol have been achieved (Lauterbach and Lenz 2019). Facultatively lithoautotrophic *R. opacus* DSM 43205 (*Nocardia opaca 1b*) offers a promising, metabolically different, alternative host for conversion of CO_2_ to chemicals (Probst and Schlegel 1973; Aggag and Schlegel 1974; Klatte et al. 1994). Unlike *C. necator* H16, it naturally accumulates fats (i.e. triacylglycerols), is Gram-positive, and contains no membrane bound hydrogen hydrogenase but only a cytoplasmic, NAD^+^-reducing hydrogenase (Aggag and Schlegel 1974; Schneider and Schlegel 1977). The cytoplasmic hydrogen hydrogenase is, however, very similar to the one of *C. necator* H16 with respect to its catalytic and molecular properties (Schneider et al. 1984). Both consist four subunits, and have FMN, nickel and iron-sulphur clusters. Like *C. necator* H16 that carries pHG1 megaplasmid with the essential genetic information for the lithoautotrophic growth, the *R. opacus* DSM 43205 hydrogen hydrogenase and Rubisco enzymes responsible for the autotrophic character are in a 270 kb linear, conjugative plasmid pHG201 (Kalkus et al. 1990; Grzeszik et al. 1997; Schwartz et al. 2003).

Heterotrophic *R. opacus* strains have been studied due to their oleaginous metabolism and versatile biodegradation pathways (Holder et al. 2011; Henson et al. 2018). Genetic engineering tools have been established for *R. opacus* (DeLorenzo et al. 2018) and used e.g. for boosting the fatty acid synthesis and for production of wax esters and alkanes (Huang et al. 2016; Lanfranconi and Alvarez 2017; Kim et al. 2019). In the present work, we engineered the autotrophic *R. opacus* DSM 43205 strain to convert CO_2_ into beta-alanine and lactic acid by expressing heterologous genes encoding aspartate decarboxylase and lactate dehydrogenase, respectively. Beta-alanine is produced by decarboxylating L-aspartate and can be used for the synthesis of different polymer precursors (i.e. acrylamide, acrylic acid, acrylonitrile and nylon-3) in the chemical industry (Könst et al. 2009; Steunenberg et al. 2013; Ko et al. 2020). Beta-alanine is also a precursor of pantothenate (vitamin B5) and coenzyme A and used as a nutritional supplement and in pharmaceutical industry as a precursor of several drugs (Khan et al. 2010; Trexler et al. 2015). As beta-alanine, also lactic acid is a polymer precursor. Several companies have already successfully commercialized the heterotrophic microbial production of lactic acid, which is not only used as a starting material for poly(lactic acid) (PLA) polymer, but also has multiple applications in chemical and other industries (Eiteman and Ramalingam 2015). Lactic acid is produced by one-step enzymatic reduction of pyruvate by lactate dehydrogenase. We report here an autotrophic production of these biopolymer precursors from CO_2_ and hydrogen gas using engineered *R. opacus* DSM 43205 strains. Two main processes have been proposed for biorefining CO_2_ into chemicals using renewable energy. In the gas fermentation process, gaseous H_2_ (generated by renewable means) is fed into a bioreactor alongside with air or oxygen and CO_2_ (Yu 2018). In the elecrobiofermentation process, only CO_2_ is fed to the bioreactor in gaseous form, while H_2_ and O_2_ are generated within the bioreactor by water electrolysis by inclusion of in-situ electrodes and renewable electricity (Liu et al. 2016). In the present work, we tested the engineered autotrophic *R. opacus* DSM 43205 strains under both process conditions. We demostrate succesful production of beta-alanine in both processes and the production of lactic acid in the gas fermentation process. We also sequenced the genome of *R. opacus* DSM 43205 and provide its improved genome assembly consisting of 18 contigs. A genome-scale metabolic model was constructed to facilitate further development of this autotrophic host for production of chemicals from CO_2_.

## Materials and methods

### Strain and plasmid construction

Synthetic genes encoding aspartate 1-decarboxylase (panD) of *Corynebacterium glutamicum* (NF003947.0) and lactate dehydrogenases of *Plasmodium falciparum* (PfLDH) (WP_074506212.1) and *Lactobacillus helveticus* (LhLDH) (WP_012211363.1) were ordered from Thermo Fisher Scientific. All three genes were optimized for expression in *R. opacus*. PDD57 and pDD65 plasmids were kindly provided by Drew M. DeLorenzo (Washington University in St. Louis, USA) (DeLorenzo et al. 2018). PDD65 is an empty plasmid with a kanamycin marker and pAL5000 (S) backbone (Ranes et al. 1990) and it was used to construct pDD57 by adding the gene encoding GFP+ expressed from a strong constitutive promoter of *Streptomyces lividans* TK24 (DeLorenzo et al. 2018). In this study, GFP+ in pDD57 was replaced with either panD, PfLDH or LhLDH by releasing GFP+ with *Nde*I + *BamH*I digestion and using the remaining plasmid backbone in Gibson assembly cloning with panD, PfLDH or LhLDH (Gibson et al. 2009). The resulting expression vectors for panD, PfLDH and LhLDH and the empty pDD65 were introduced into *R. opacus* cells by electroporation. For electroporation, *R. opacus* was grown in 50 ml of TSB media overnight at 30°C and cells were harvested by centrifugation and washed twice with cold 20 mM HEPES, 15% glycerol pH 7.2 and once with 5 mM HEPES, 15% glycerol pH 7.2. Cells were suspended into approximately 800 µl of 5 mM HEPES, 15% glycerol pH 7.2 and 80 µL of suspension was mixed with 1 µg of plasmid DNA. Electroporation was carried out in 1 mm electroporation cuvettes with the following settings: 25 µF, 400 Ω, 2.5 kV. After the electroporation, cells were incubated at 30°C for 3 h in 800 µL of SOC media before plating them on TSA plates containing 50 µg mL^-1^ kanamycin. The presence of expression vectors in the resulting transformants ROP-pDD65 (ctrl), ROP-PfLDH, ROP-LhLDH and ROP-panD was verified by colony PCR by using the plasmid specific DNA oligomers and DreamTaq DNA polymerase (Thermo Fisher Scientific).

### Growth media

All cultivations were grown on a modified DSM-81 mineral medium (www.dsmz.de/microorganisms/medium/pdf/DSMZ_Medium81.pdf). The major chloride salts in the original recipe were replaced with corresponding sulfates to reduce the formation of chlorine gas during the electrolysis. Additionally, the vitamins were omitted from the recipe. In our trials, addition of the vitamin solution into media did not enhance the growth (data not shown). Furthermore, beta-alanine is a precursor of one of the B vitamins (pantothenate) and therefore the presence of pantothenate in media could interfere with the beta-alanine metabolism. The final medium composition per one liter was: 2.3 g KH_2_PO_4_, 2.9 g Na_2_HPO_4_·2H_2_O, 5.45 g Na_2_SO_4_, 1.19 g (NH_4_)_2_SO_4_, 0.5 g MgSO_4_·7H_2_O, 11.7 mg CaSO_4_·2H_2_O, 4.4 mg MnSO_4_·H_2_O, 5 mg NaVO_3_, 0.5 g NaHCO_3_, 5 mg ferric ammonium citrate, 0.5 mg ZnSO_4_·7H_2_O, 1.5 mg H_3_BO_3_, 1 mg CoCl_2_·6H_2_O, 50 μg CuCl_2_·2H_2_O, 0.1 mg NiCl_2_·6H_2_O, 0.15 mg Na_2_MoO_4_·2H_2_O. 50 µg mL^-1^ kanamycin and/or 20 g L^-1^ glucose were added into media when appropriate.

### Inocula and shake flask cultivations

*R. opacus* transformants were maintained on TSA plates containing 50 µg ml^-1^ kanamycin. Inocula for glucose cultivations were grown at 30 °C in 10 mL of modified DSM-81 media with 20 g L^-1^ glucose and 50 µg mL^-1^ kanamycin in 50 mL erlenmayer flasks. Glucose shake flask cultivations were carried out in 250 mL erlenmayer flasks. Cultivations were started from optical density (at 600 nm) (OD_600nm_) of 0.1 and carried out at 30 °C with 220 rpm shaking in 50 mL of the same media that was used to grow the inocula. CO_2_ shake flask cultivations or inocula for electrobiofermentations were started by inoculating a loopfull of cells directly from TSA plate into 20 mL of modified DSM-81 media with 50 µg mL^-1^ kanamycin. 100 mL erlenmayer flask were incubated in a sealed container with 130 rpm shaking and fed with 20 mL min^-1^ air, 4 ml min^-1^ H_2_ and 8 mL min^-1^ CO_2_. The resulting gas composition in a container was 49% N_2_, 25% CO_2_, 13% H_2_ and 13% O_2_.

### Electrobiofermentations

Electrobiofermentations were performed in 100 ml MR-1194 Bulk Electrolysis cell vials (BASi, West Lafayette, IN, USA) with custom-made Teflon lids. The electrobiofermentation setup was similar than the one used by Marsili et al. (2008) (Marsili et al. 2008). 70 mL of modified DSM-81 media supplemented with 13 µL of Componenta VO antifoam (Ecolab, Netherland) was inoculated with preculture to obtain the initial OD_600_ 0.2. A gas mix consisting of 20% CO_2_ and 80% N_2_ (AGA, Espoo, Finland) was fed at 6 mL min^-1^ rate. The gas feed rate was controlled by using a mass flow module (Medicel, Espoo, Finland). The gas was humified with water by bubbling the input gas stream through a sterile filter (Whatman, 0.2 µm, Sigma-Aldrich, Saint Louis, MO, USA) into sterilized water before feeding into the reactor. The reactor temperature was maintained at 30 °C with water circulated through the heating jacket of the reactor using two external water baths (Julabo, Seelbach, Germany and VWR International, Radnor, PA, USA). Liquid in the reactor was agitated with a magnetic stirrer bar rotating at 400 rpm. Coiled titanium wire coated with a thin layer of iridium oxide (IrO) was used as the anode (∅ 1.5 mm, Magneto Special Anodes, Schiedam, Netherlands) and a coiled stainless steel (SS) capillary was used as the cathode (∅ 1.6 mm, 316L-SS, Pfeiffer Vacuum GmbH, Asslar, Germany). The electrode surface areas were 0.15 cm^2^ and 12.3 cm^2^ for Pt and IrO electrodes, respectively. The voltage and current were controlled using a Wavenow potentiostat (Pine Research Instrumentation, Grove City, PA, USA), and the AfterMath computer program (Version 1.3.7060, Pine Research Instrumentation, Grove City, PA, USA). The electrolysis was performed at a current of 18 mA (chronopotentiometry).

### Analytical methods

Biomass growth was measured from cultivations by taking 1-2 mL samples and measuring OD_600nm_ using a UV-1201, UV-vis spectrophotometer (Shimadzu, Kyoto, Japan). Samples of high optical density were diluted to obtain sample ODs in the range of 0.1 – 0.3. Cell dry weight (CDW) was measured from 2 ml cultivation samples by separating the cells by centrifugation and washing them twice with MilliQ water before drying them at 105 °C overnight. Alternatively, OD_600nm_ of culture broth was converted to dry cell weight (CDW) per L^-1^ culture broth by using a standard curve prepared previously.

Extracellular concentrations of glucose and lactic acid from glucose shake flask cultivations were analysed by high-performance liquid chromatography (HPLC) on an Fast acid and Aminex HPX-87H columns (BioRad Laboratories, Hercules, CA) with 2.5 mM H_2_SO_4_ as eluant and a flow rate of 0.5 mL min^-1^. The column was maintained at 55°C. Peaks were detected using a Waters 410 differential refractometer and a Waters 2487 dual wavelength UV (210 nm) detector. Extracellular concentration of lactic acid from CO_2_-cultivations was analysed by GC-MS. Each cell culture supernatant sample (50 μL) was spiked with internal standard (10 μL of 3-hydroxybutyric acid-1,3-13C2 acid) and the sample was evaporated into dryness under N_2_ flow. The dried residues were derivatized with a mixture of 50 µL of pyridine and 50 µL of N-Methyl-N-(trimethylsilyl) trifluoroacetamide (MSTFA) reagent containing 1% of trimethylchlorosilane (TMCS) as a catalyst (70°C, 60 min).

The sample compositions were analysed on an Agilent 6890 gas chromatograph (GC) combined with Agilent 5973 mass selective detector (MSD). The injector (injection volume 1 µL) and inlet temperature was 250 °C, and the oven temperature was increased from 50 °C to 310 °C. The analyses were performed on an Agilent DB-5MS capillary column (30 m, ID 250 µm, film thickness 0.25 µm; Agilent 122-5532). Lactic acid was quantified by monitoring its m/z ion ratio of 191. The calibration range for lactic acid was 0.3 – 33 µg per sample and the quantitation ion was m/z 191.

Extracellular beta-alanine concentration was analysed by ultra-performance liquid chromatography (UPLC). 250 µL of cell culture supernatant was deproteinized by adding 750 µL of ethanol (99.5 %), the samples were mixed and centrifuged. The supernatant was transferred to a new vial and the samples were concentrated under a stream of N_2_. Finally, the volume was adjusted to 80 µL, 20 µL of borate buffer was added and 10 µl of the sample solution were taken for the analysis. The internal standard solution (norvaline, 25 μM), MassTrak™ AAA borate buffer and 6-aminoquinolyl-N-hydroxysuccinimidyl carbamate reagent were added, and sample mixture was instantly vortexed before incubation at 55 °C for 10 min. Amino acid standard mixtures were derivatized identically with the samples.

UPLC analysis was performed using an Acquity UPLC system, Waters (Milford, MA, USA) with a Waters UV detector. Chromatography was performed using an Acquity MassTrak™ (2.1 × 100 mm, 1.7 μm) column, Waters (Milford, USA), kept at 43 °C. Injection volume was 1 µL. Separation was performed using gradient elution with 10% (v/v) MassTrak™ AAA eluent A concentrate in water (A) and MassTrak™ AAA eluent B (B) at a flow rate of 0.4 mL min^-1^ using a gradient elution program. Signal for beta-alanine was detected at 260 nm. MassTrak™ Amino Acid Analysis (AAA) derivatization kit, Mass TRAK™ Amino Acid Analysis concentrate A and eluent B were obtained from Waters (Milford, MA, USA). Amino Acid Standard Solution, Amino Acid Standards Physiological, Basics, L-isoleusine, glutamine and norvaline were obtained from Sigma-Aldrich (St. Louis, Missouri, USA).

### Next generation sequencing

Sequencing libraries were prepared by BaseClear BV (Leiden, The Netherlands), using Illumina Nextera XT kit for short-read sequencing and 10KB PacBio library preparation technique for long-read sequencing. BaseClear BV (Leiden, The Netherlands) performed short-read Illumina-based paired-end sequencing (HiSeq 2500, 2×125bp) at a depth of 200MB, and long-read sequencing using PacBio Sequel SMRT platform. BaseClear BV (Leiden, The Netherlands) performed the quality filtering and delivered the resulting filtered raw data of the sequence reads. We used FastQC (version 0.11.7) for analysing the quality of raw sequencing reads to confirm that the short-read data showed high base call quality across all the bases. As the PacBio Sequel platform does not report the quality values for the base calls, the long-read data was excluded from the post-hoc FastQC-based quality analysis.

### Genome assembly and annotation

*De novo* genome assembly was performed using Unicycler v0.4.6 (Wick et al. 2017), utilizing the combination of the PacBio-derived long reads and Illumina-derived short reads. Gene prediction was performed using the bacterial genome annotation pipeline Prokka v1.14.5 (Seemann 2014), and functional annotations were performed using eggNOG (Huerta-Cepas et al. 2016) and Pannzer (Törönen et al. 2018).

### Comparison with other Knallgas bacteria

In order to compare the metabolic capabilities of the *Rhodococcus opacus* DSM 43205 genome assembly and the functional annotations derived in this study, we compiled a list of Knallgas strains (n = 68) based on literature evidence. Out of these 68, we were able to retrieve annotated genome assemblies of 50 strains from NCBI GenBank and RefSeq databases. In addition, we included the genome assembly of *N. nitrophenolicus* strain KGS27 (unpublished) in the comparison. The collected genomes were verified by the presence of RuBisCo (EC: 4.1.1.39) and at least one of the hydrogenases (EC: 1.12.1.2, EC: 1.12.1.3 and EC:1.12.5.1) or by empirical literature evidence for whether the strain was shown to grow on CO_2_ alone as the carbon source. We also included the *R. opacus* strains 1CP and B4 for reference even though they are not Knallgas bacteria. Based on these selection criteria, we included 45 strains in the comparison (Supplementary Table 1). The list of enzymes (EC numbers) in each KEGG pathway was derived from the KEGG REST API (https://www.kegg.jp/kegg/rest/keggapi.html) and those pathways whose 50% of the enzymes are present in the *Rhodococcus opacus* DSM43205 assembly were used in clustering of selected strains.

**Table 1.** Volumetric and specific rates of lactic acid and beta-alanine production from glucose and CO_2_ in shake flask and electrobiofermetation cultivations. Lactic acid was not detected (nd) from electrobiofermentations.

|  | Glucose shake flask |  |  |  | CO<sub>2</sub> shake flask |  |  | CO<sub>2</sub> electrobiofermentation |  |  |
| --- | --- | --- | --- | --- | --- | --- | --- | --- | --- | --- |
|  | mg l<sup>-1</sup><br>h<sup>-1</sup> | mg g<sup>-1</sup><br>glucose | mg g<sup>-1</sup><br>CDW | mg g<sup>-1</sup><br>CDW h<sup>-1</sup> | mg l<sup>-1</sup><br>h<sup>-1</sup> | mg g<sup>-1</sup><br>CDW | mg g<sup>-1</sup><br>CDWh<sup>-1</sup> | mg l<sup>-1</sup><br>h<sup>-1</sup> | mg g<sup>-1</sup><br>CDW | mg g<sup>-1</sup><br>CDW h<sup>-1</sup> |
| Strain | Lactic acid |  |  |  |  |  |  |  |  |  |
| ROP-PfLDH | 31.2 | 129.6 | 434.6 | 19.8 | 0.63 | 151.8 | 0.7 | nd | nd | nd |
| ROP-LhLDH | 25.5 | 116.4 | 369.7 | 16.8 | 0.37 | 66.5 | 0.4 | nd | nd | nd |
|  | Beta-alanine |  |  |  |  |  |  |  |  |  |
| ROP-panD | 0.80 | 1.62 | 0.42 | 0.01 | 0.01 | 1.92 | 0.01 | 0.0016 | 0.1632 | 0.0005 |

### Genome-scale metabolic model reconstruction and simulations

A genome-scale metabolic model for *R. opacus* DSM 43205 was reconstructed using CarveMe (Machado et al. 2018). The bacterial universal metabolic model constructed based on the reactions obtained from BiGG models database (King et al. 2016) was used as the reference. The proteome sequence of *R. opacus* DSM 43205 derived in the gene prediction step was used to calculate the reaction scores for the model reconstruction as follows. The *R. opacus* DSM 43205 protein sequences were aligned to the sequences of the BiGG genes using Diamond (Buchfink et al. 2014) and the best alignment score for each BiGG gene was used as gene-level score. The gene-level scores were converted to reaction-level scores via the gene-protein-reaction (GPR) rules as follows. The protein-level scores were calculated as the average gene-level score of all subunits in each protein complex, and the maximum of the protein-level scores of all isozymes that catalyze each reaction was used as the reaction-level score. The reaction-level scores were normalized with the median reaction score. Enzyme-catalyzed reactions without genetic evidence were given score = –1 and the spontaneous reactions were assigned score = 0. The *R. opacus* DSM 43205 genome-scale metabolic model simulations were performed with *framed* (https://github.com/cdanielmachado/framed), a metabolic modelling package for python, using IBM ILOG CPLEX LP-solver v. 12.8.0 function *cplexlp*.

## Results

### Heterotrophic and autotrophic lactic acid production

Genes encoding lactate dehydrogenases of *P. falciparum* (PfLDH) and *L. helveticus* (LhLDH) were cloned into pDD57 expression plasmid under constitutive promoter of *Streptomyces lividans* TK24 and transformed into *R. opacus* DSM 43205. First, the production of lactic acid by three ROP-PfLDH and three ROP-LhLDH transformants was studied in shake flask cultures on 20 g L^-1^ glucose. At maximum, 742 mg L^-1^ lactic acid was produced by ROP-PfLDH at a rate of 31 mg L^-1^ h^-1^ and a specific productivity of 20 mg g CDW^-1^ h^-1^. ROP-LhLDH produced lactic acid at maximum 608 mg L^-1^ at a rate of 26 mg L^-1^ h^-1^ and a specific productivity of 17 mg g CDW^-1^ h^-1^ (Fig. 1a & Table 1). In both cases, the lactic acid accumulation took place at the beginning on the cultivation during the 20 h phase when both glucose consumption and biomass accumulation were slow. After that, during the following 10 h, cells consumed the produced lactic acid and almost all glucose rapidly resulting in biomass accumulation that continued until 53 h up to an OD_600nm_ of 35. The pH in the cultivations of the lactic acid producing strains were lower compared with the control strain with the empty expression vector during the lactic acid production phase but did not remarkably differ from the control at the later stages of cultivations (Fig. 1b). Interestingly, the lactic acid producing strains consumed glucose slightly faster and reached a slightly higher biomass than the control strain.

**Figure 1.**
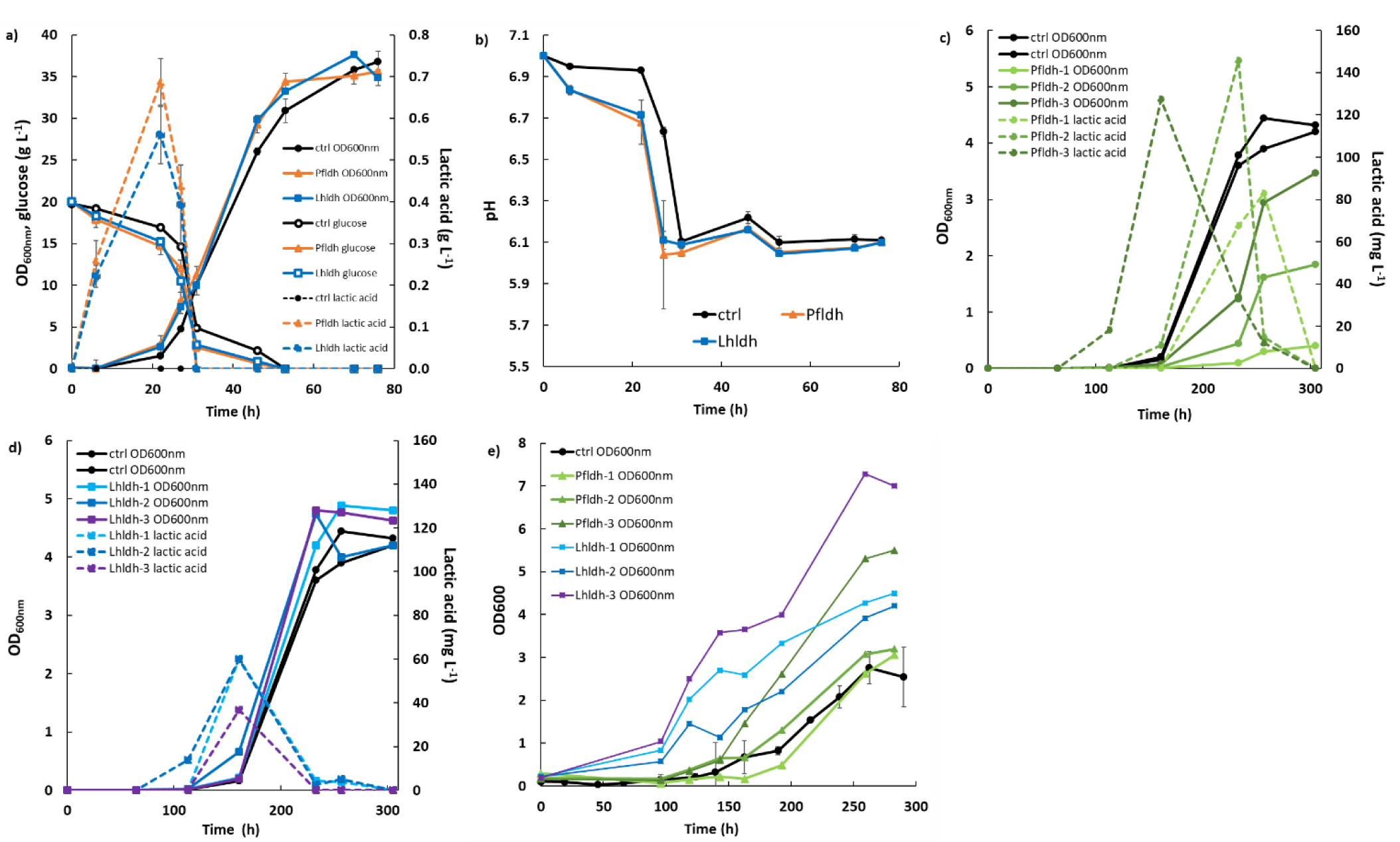
Heterotrophic and autotrophic lactic acid and biomass production by *R. opacus* DSM 43205 transformants expressing lactate dehydrogenase of *L. helveticus* (LhLDH) or *P. falciparum* (PfLDH). a) Glucose consumption, lactic acid (g L^-1^) and biomass production (OD_600nm_) from 20 g L^-1^ glucose in shake flask cultures, b) pH of glucose cultivations, c) lactic acid (mg L^-1^) and biomass production of LhLDH expressing strains in shake flask cultures on CO_2_, d) lactic acid (mg L^-1^) and biomass production of PfLDH expressing strains in shake flask cultures on CO_2_, e) biomass production in electrobiofermentations from CO_2_.

Next, lactic acid production by ROP-PfLDH and ROP-LhLDH transformants was studied under autotrophic conditions in shake flask cultures incubated in a sealed container under a gas with the composition of 49% N_2_, 25% CO_2_, 13% H_2_ and 13% O_2_. The growth of the control strains and ROP-PfLDH and ROP-LhLDH initiated after ca. 100 h lag phase and the *R. opacus* strains ROP-PfLDH and ROP-LhLDH produced lactic acid at maximum concentrations of 146 mg L^-1^ and 61 mg L^-1^, respectively (Fig. 1c and d). The specific lactic acid productivities of both *R. opacus* strains ROP-PfLDH (0.7 mg g CDW^-1^ h^-1^) and ROP-LhLDH (0.4 mg g CDW^-1^ h^-1^) were significantly lower than in the glucose cultivation (Table 1.). Similar to glucose cultivations, the peak lactic acid production coincided with the early phases of growth. In contrast to glucose cultivations, where there was no significant difference in growth and lactic acid production between the *R. opacus* strains ROP-PfLDH and ROP-LhLDH, from CO_2_ the ROP-LhLDH produced less lactic acid and more biomass while ROP-PfLDH transformants accumulated less biomass and produced over two fold more lactic acid. Lactic acid production was also studied in electrobiofermentations that were sparged with 20% CO_2_. H_2_ and O_2_ were provided by water electrolysis that was performed at a constant current of 18 mA. Under these conditions, both ROP-PfLDH and ROP-LhLDH accumulated biomass but no lactic acid production was detected (Fig. 1e). All ROP-LhLDH transformants accumulated more biomass than the control strain similar to shake flask cultures that were grown with CO_2_ as carbon source.

### Heterotrophic and autotrophic beta-alanine production

Gene encoding aspartate 1-decarboxylase (panD) of *C. glutamicum* was cloned into the pDD57 expression plasmid under constitutive promoter of from *Streptomyces lividans* TK24 and transformed into *R. opacus* DSM 43205. On glucose ROP-panD transformants and the control strain containing the empty pDD65 grew almost equally well (Fig. 2a) up to OD_600nm_ 38 and the *R. opacus* strain ROP-panD produced at maximum 25 mg L^-1^ extracellular beta-alanine with specific productivity of 0.01 mg g CDW^-1^ h^-1^ while the control strain produced 1.6 mg L^-1^. In both cases, beta-alanine production took place at the early growth phase when glucose was still consumed slowly. Thereafter, the produced beta-alanine was rapidly consumed during the exponential growth phase. In shake flasks in a sealed container under a gas atmosphere containing 49% N_2_, 25% CO_2_, 13% H_2_ and 13% O_2_ and when CO_2_ was used as a carbon source maximum 1.8 mg L^-1^ beta-alanine was produced (Fig. 2b). The specific productivity was the same as in glucose cultivation (Table 1.). Also, from CO_2_ beta-alanine was produced during the ca. 100 h lag phase before the growth initiated and it was then consumed during the growth. Beta-alanine producing transformants reached 17% higher OD_600nm_ values than the control strains (OD_600nm_ 5 vs. 4.2). Beta-alanine production was also demonstrated in electrobiofermentations sparged with 20% CO_2_ and with provision of H_2_ and O_2_ by water electrolysis at a constant current of 18 mA. Under these conditions, ROP-panD strains produced 345 µg L^-1^ beta-alanine from CO_2_ (Fig. 2c). Beta-alanine started to accumulate already during the lag phase of the cultivations, but the highest production coincided with the growth phase of the cells. After that, the produced beta-alanine was consumed from the media. ROP-panD strains reached 50% higher biomass (OD_600nm_ 5.5) than the control strains.

**Figure 2.**
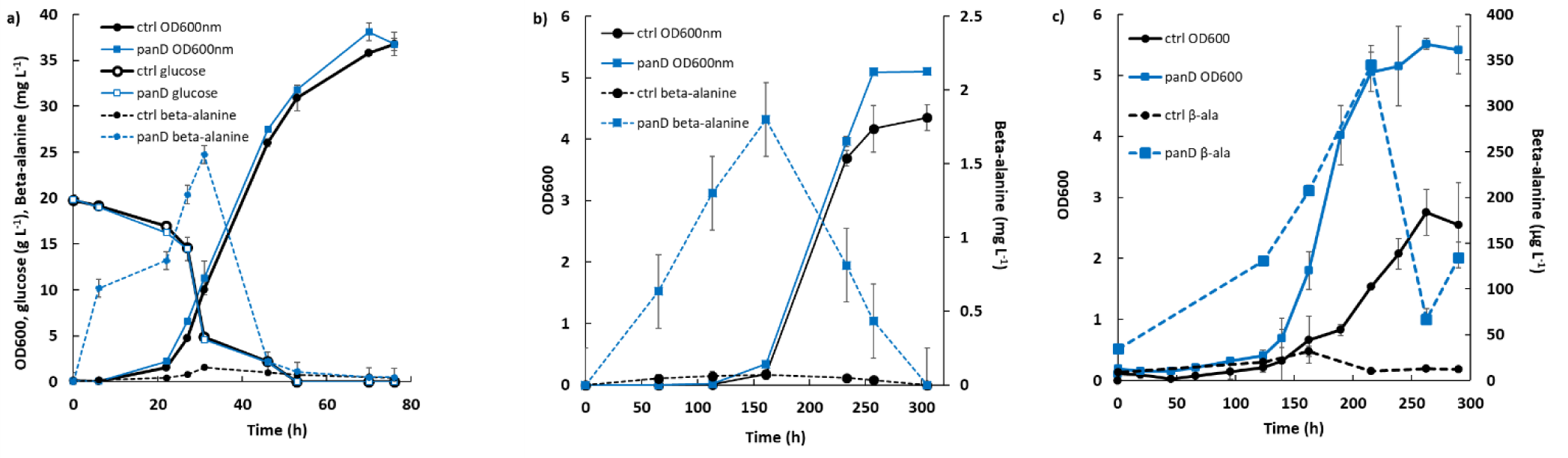
Heterotrophic and autotrophic beta-alanine and biomass production by *R. opacus* DSM 43205 transformants expressing aspartate decarboxylase of *C. glutamicum* (panD). a) Glucose consumption, beta-alanine (mg L^-1^) and biomass production (OD_600nm_) from 20 g L^-1^ glucose in shake flask cultures, b) beta-alanine (mg L^-1^) and biomass production of panD expressing strain in shake flask cultures on CO_2_, c) beta-alanine (µg L^-1^) and biomass production of panD expressing strains in electrobiofermentations from CO_2_.

### Genome analysis of *R. opacus* DSM 43205

The Illumina data set contained over 5.53 million paired-end reads, with lengths that ranged from 50 to 126. The PacBio-based long read data set contained over 758 thousand reads with lengths that ranged from 50 to 41,685. The Unicycler-based genome assembly contained eighteen contigs (length > 1500 bases) together representing a genome of 8,942,682 bases. The largest contig (and N50) was 6,484,583 bases long. The GC content of the assembly was 67%. A number of (n = 8418) gene coding sequences (CDS) were found in the genome assembly. Of these, majority of the CDS (n = 5999) were found on the largest contig. A number of the sequences (n = 2684) were identified as metabolic enzymes with known EC numbers, which mapped to over 145 metabolic pathways in the Kyoto Encyclopedia of Genes and Genomes (KEGG) database. The autotrophic marker ribulose-1,5-bisphosphate carboxylase (RuBisCo, EC:4.1.1.39) was found in the genome assembly (on the 5^th^ largest contig) confirming the presence of Calvin cycle, along with the hydrogenase (NAD-reducing hydrogenase HoxS subunits, EC: 1.12.1.2 in contigs 2, and 5) (Supplementary Figure 1). Together, these markers confirm the experimentally observed CO_2_ metabolism of the *R. opacus* DSM 43205 strain.

### Comparison of R. opacus DSM 43205 with other Knallgas bacteria

In comparison of *R. opacus* DSM 43205 genome with other Knallgas bacteria, those KEGG pathways whose 50% of the enzymes are present in the *Rhodococcus opacus* DSM43205 assembly are shown in Figure 3a. In addition, a comparison of the Knallgas bacterial strains has also been provided in terms of a list of key enzymes related to the Knallgas activity including RuBisCo, H_2_ hydrogenases, CO_2_ transporters, and Calvin-Benson-Bassham cycle enzymes (Fig. 1b).

**Figure 3.**
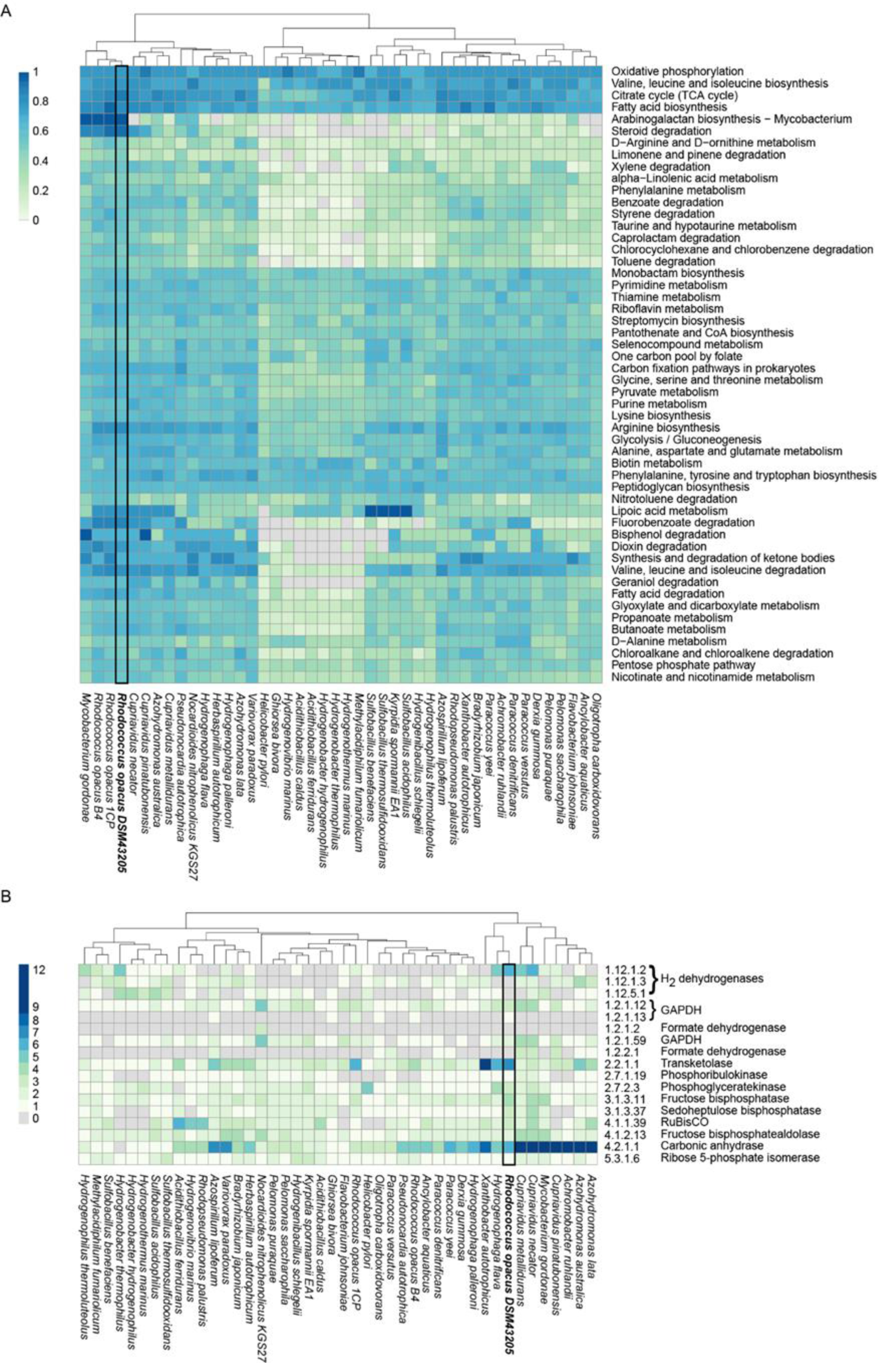
Metabolic pathways and key enzymes in the Knallgas bacteria and *R. opacus* 1CP and B4 strains. (a) A clustering of Knallgas strains based on the proportion of the enzymes found in each metabolic pathway (0 in a cell represents that none of the enzymes are found, and 1 represents that 100% of the enzymes are found). Only the pathways where *R. opacus* DSM 43205 has a minimum of 50% of the enzymes are shown. (b) A clustering of Knallgas strains based on the number of coding sequences found in each genome for a key set of enzymes related to CO_2_ metabolism.

When clustering Knallgas bacteria in terms of the percentage of metabolic pathway enzymes found, the strains of *R. opacus* clustered together (Fig. 1a). Majority of the strains are enriched with the enzymes in oxidative phosphorylation, fatty acid biosynthesis, TCA cycle, and biosynthesis of the essential amino acids, valine, leucine and isoleucine. The cluster containing the *R. opacus* strains was more enriched for enzymes in arabinogalactan biosynthesis and steroid degradation than others.

When clustering Knallgas bacteria in terms of the number of sequences encoding for a set of key enzymes required for CO_2_ metabolism (namely, RuBisCo, H_2_ hydrogenases, CO_2_ transporters, and Calvin-Benson-Bassham cycle enzymes), *R. opacus* DSM 43205 clustered more closely with *Hydrogenaphaga flava, Xanthobacter autotrophicus* and some strains of *Cupriavidus* than with other *R. opacus* strains (Fig. 1b). The cluster containing the *R. opacus* DSM 43205 was relatively more enriched for the hydrogenase EC: 1.12.1.2 and transketolase EC: 2.2.1.1. The cluster containing the *R. opacus* DSM 43205 and *C. necator* was the most enriched with sequences coding for carbonic anhydrase (EC: 4.2.1.1).

### Genome-scale metabolic model simulations

Automatically reconstructed genome-scale metabolic model of *R. opacus* DSM 43205 (containing 2267 reactions and 1499 metabolites) was manually curated for aerobic autotrophic growth with soluble hydrogen hydrogenase with oxygen as the final electron acceptor (Supplementary material). The model-predicted that the hydrogen to carbon dioxide utilization ratio for optimal growth was 4.11 mol H_2_ per mol CO_2_. At higher ratio, the growth was predicted to be limited by the availability of CO_2_, whereas at lower utilization ratio the growth was predicted to be H_2_-limited. Model simulations were performed also for predictions on lactic acid and beta-alanine production using *R. opacus* DSM 43205. The H_2_ to CO_2_ utilization ratios for the optimal synthesis of these compounds were predicted with model simulations very similar for the two products and slightly lower than for growth. The predicted utilization ratios were 3.07 mol H_2_ per mol CO_2_ and 3.19 mol H_2_ per mol CO_2_ for lactic acid and beta-alanine, respectively.

## Discussion

*R. opacus* strain DSM 43205 is an interesting representative of less known aerobic facultative chemolithotrophs that potentially have the capability for CO_2_-based chemical synthesis (Aggag and Schlegel 1974; Brigham 2019). In the present study, CO_2_ or glucose were converted to the biopolymer precursor beta-alanine or lactic acid by *R. opacus* DSM 43205 by the action of heterologous aspartate decarboxylase and lactate dehydrogenase, respectively. Beta-alanine production from CO_2_ was demonstrated also in electrobiofermentations where H_2_ and O_2_ were provided by water electrolysis. Use of H_2_ as a gaseous substrate in fermentations can be difficult due to its low solubility and safety issues. The *in-situ* water electrolysis is therefore a sustainable and safer method for hydrogen provision.

Beta-alanine is a precursor of pantohenate and CoA biosynthesis and its biosynthesis involves at least three enzymatic reactions from phosphoenolpyruvate depending on whether aspartate is formed directly from oxaloacetate or through the TCA cycle (Piao et al. 2019). The *R. opacus* genome encodes genes for both of these routes. It also possesses an endogenous gene for aspartate decarboxylase and a small amount of beta-alanine was produced in both glucose and CO_2_ cultivations with the control strain. Overexpression of panD from *C. glutamicum* increased the beta-alanine production significantly from both carbon sources. Interestingly, ROP-panD strains produced also more biomass than the control from both glucose and CO_2_, especially in electrobiofermentations. There is no evident reason for this but possibly consumption of the produced beta-alanine from the growth media boosted the carbon metabolism and growth.

*R. opacus* does not natively produce lactic acid in the single step enzymatic reduction of pyruvate, which is an abundant, central metabolite of glycolysis. Here, we overexpressed in *R. opacus* lactate dehydrogenase genes from *L. helveticus* and *P. falciparum* with different catalytic properties. PfLDH has a significantly lower K_m_ for pyruvate than LhLDH (0.03 mM vs 0.25 mM) and its catalytic efficiency is significantly higher (Novy et al. 2017). ROP-PfLDH strains produced almost 2.5-fold more lactic acid from CO_2_ in shake flask cultivations than ROP-LhLDH strains. This may be attributed to the lower K_m_ of PfLDH for pyruvate. The lower biomass production by ROP-PfLDH strains was possibly due to channeling of more carbon to lactic acid, although it is likely that there are also other reasons for the difference of produced biomass between ROP PfLDH and LhLDH strains because the concentration of produced lactic acid was so small compared to biomass difference. Interestingly, both strains produced almost equal amount of lactic acid from glucose, which is possibly due to a higher concentration of available pyruvate under these conditions.

Lactic acid was not produced into growth media in electrobiofermentations. It is possible that in small electrobioreactors water splitting with relatively low current resulted in limited generation of H_2_ that restricted the NADH provision by the H_2_ hydrogenase. The actual redox requirements for lactic acid and beta-alanine synthesis are equal and theoretically optimal yields of the products would require very similar H_2_ to CO_2_ utilization ratios (predicted by the genome-scale metabolic model simulations). However, lactic acid synthesis may become thermodynamically unfavorable already at smaller decrease in NADH/NAD^+^ ratio than beta-alanine synthesis. In these cultivations, especially ROP-LhLDH strains grew to higher cell densities than control and ROP-PfLDH strains and the reasons for this are currently unknown.

Despite the difference in length of lactic acid and beta-alanine biosynthetic routes from pyruvate, the production of both compounds was observed in both cases already at lag phase and at the early phase of exponential growth independent on whether glucose or CO_2_ was available as the carbon source. Especially, the lactic acid production appeared to increase the initial glucose consumption rate in the beginning of the cultivations. After that, both lactic acid and beta-alanine were consumed. In case any lactic acid or beta-alanine production took place during the exponential growth phase, both compounds were immediately consumed and could not be detected in the supernatant.

Simple overexpression of aspartate decarboxylase and lactate dehydrogenases in *R. opacus* allowed us to demonstrate the production of beta-alanine and lactic acid from CO_2_ but economically viable production titers would clearly require more elaborate metabolic engineering efforts. As an example, in *Escherichia coli*, efficient production of beta-alanine from glucose requires overexpression of all genes encoding enzymes of the reductive branch of TCA cycle and deletion of several pathways for side products (Piao et al. 2019; Zou et al. 2020). In *E. coli*, the uptake of beta-alanine is performed by an active amino acid transporter (Schneider et al. 2004). Deletion of the corresponding transporter from *R. opacus* could possibly prevent the beta-alanine utilization from growth media. Likewise, further enhanced lactic acid production would require more extensive metabolic engineering as carried out for *E. coli* (Jiang et al. 2017).

In order to improve *R. opacus* DSM 43025 and its industrial applicability for chemical production by metabolic engineering, it would be important to increase the understanding of the characteristics of its heterotrophic and lithoautotrophic metabolism and optimize the entire metabolic network at the systems level. In *R. opacus* DSM 43025, carbon from both glucose and CO_2_ was predominantly assimilated into biomass. Using a combination of PacBio-based long-read and Illumina-based short-read next-generation sequencing data, we provided an improved assembly of *R. opacus* DSM 43205 in 18 contigs, in comparison to the existing 382-contig assembly (https://www.ncbi.nlm.nih.gov/assembly/GCF_001646735.1/). The improved genome assembly of *R. opacus* DSM43025 together with the genome-scale metabolic model curated for aerobic autotrophic metabolism will promote the discovery of further engineering strategies for *R. opacus* DSM43025.

## Conclusions

Microbial CO_2_ fixation using solar power generated sustainable H_2_ as an energy source has the potential to become a major strategy to use CO_2_ as a raw material. Conversion of CO_2_ to chemicals in bioprocesses by lithoauthotrophic, H_2_-oxidizing bacteria is a promising way to reduce global CO_2_ emissions.

The novel hosts resulting from the work contribute to the transition from CO_2_-releasing manufacture of chemicals to CO_2_-fixing bioprocesses. The molecular biology tools, cultivation methods, and metabolic models developed in this project will facilitate further studies of still mostly unexplored lithoautotrophic microbial species.

## Acknowledgements

This work was financially supported through Academy of Finland research grants MOPED (Decision No. 295883) and Optobio (Decision No. 287011), Business Finland research grant Fermatra (Diary No. 908/31/2016) and a joint research grant by the Technology Industries of Finland Centennial Foundation and the Jane and Aatos Erkko Foundation on “Feed and food from CO_2_ and electricity – the research and piloting of future protein production”. Paula Jouhten would like to acknowledge funding from the Academy of Finland (decision no. 310514).

