## Supplemental Fig. 1 and Table 1 for "Production of biopolymer precursors beta-alanine and L-lactic acid from CO_2_ with metabolically versatile *Rhodococcus opacus* DSM 43205"

Supplementary Figure 1: RuBiSCo and hydrogenases found in the *Rhodococcus opacus* DSM 43205 assembly confirming the hydrogen oxidizing autotrophic (HOA) activity. The largest seven contigs are shown in the picture with markings of where RuBiSCo (green color labels) and hydrogenases (red color labels) were found. Note that the NAD-reducing hydrogenase HoxS subunits (EC: 1.12.1.2) found on contigs 2 and 5 are relevant for HOA activity, but not the periplasmic NiFeSe hydrogenaes subunits found on contig 1.

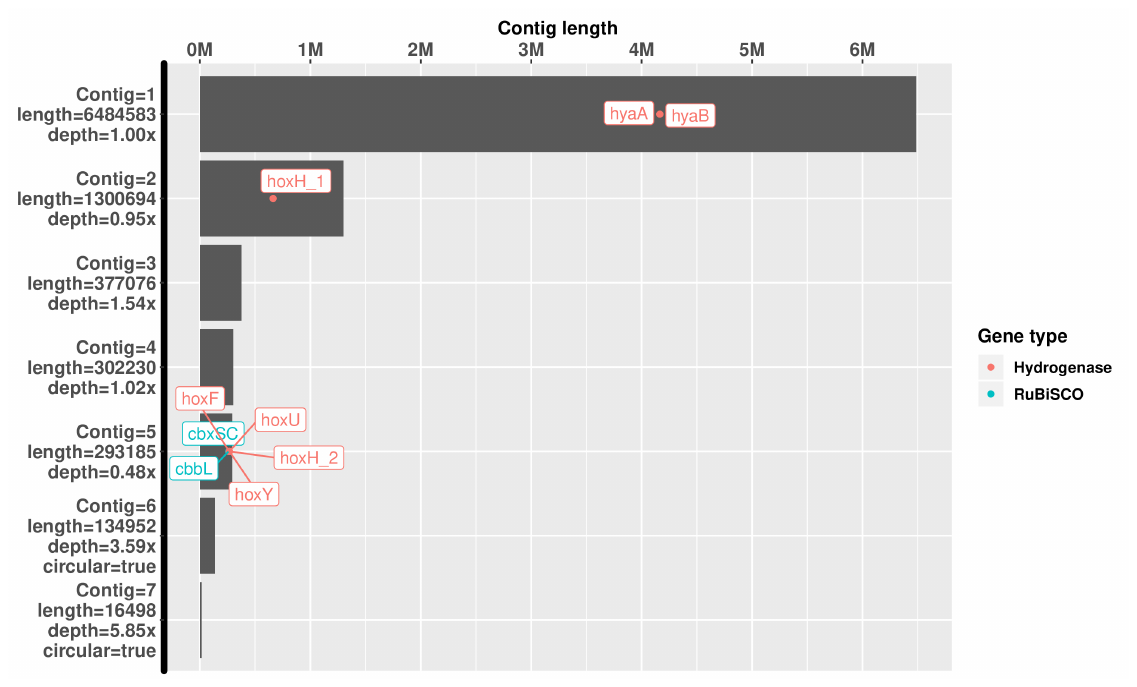

Supplementary Table 1: The list of Knallgas bacteria included in the comparison with *Rhodococcus opacus* DSM 43205.

| **Species** | **Strain** |
| --- | --- |
| Achromobacter ruhlandii | 6241 |
| Acidimicrobium ferrooxidans | DSM 10331 |
| Acidothiobacillus caldus | ATCC 51756 |
| Acidothiobacillus ferridurans | JCM 18981 |
| Acidovorax delafieldii | 2AN |
| Ancylobacter aquaticus | UV5 |
| Azohydromonas australica | DSM 1124 |
| Azohydromonas lata | NBRC 102462 |
| Azorhibozium caulinodans | ORS 571 |
| Azospirillum lipoferum | 4B |
| Bradyrhizobium japonicum | CCBAU 15354 |
| Cupriavidus metallidurans | CH34 |
| Cupriavidus necator | H16 |
| Cupriavidus pinatubonensis | DSM 19553 |
| Derxia gummosa | DSM 723 |
| Ghiorsea bivora | TAG-1 |
| Herbaspirillum autotrophicum | IAM 14942 |
| Hydrogenibacillus schlegelii | DSM 2000 |
| Hydrogenobacter hydrogenophilus | DSM 2913 |
| Hydrogenobacter thermophilus | TK-6 |
| Hydrogenophaga flava | NBRC 102514 |
| Hydrogenophaga palleroni | NBRC 102513 |
| Hydrogenophilus thermoluteolus | TH-1 |
| Hydrogenothermus marinus | VM1 |
| Hydrogenovibrio marinus | MH-110 |
| Kyrpidia spormannii | EA-1 |
| Methylacidiphilum fumariolicum | SolV |
| Mycobacterium gordonae | CTRI 14-8773 |
| Nocardioides nitrophenolicus KGS27 | KGS27 |
| Oligotropha carboxidovorans | ATCC 49405 |
| Paracoccus denitrificans | PD1222 |
| Paracoccus versutus | DSM 582 |
| Paracoccus yeei | ATCC BAA-599 |
| Pelomonas puraquae | CCUG 52769 |
| Pelomonas saccharophila | DSM 654 |
| Pseudonocardia autotrophica | NRRL B-16064 |
| Rhodocuccus opacus DSM 43205 | DSM 43205 |
| Rhodocuccus opacus 1CP | 1CP |
| Rhodocuccus opacus B4 | B4 |
| Rhodopseudomonas palustris | HaA2 |
| Sulfobacillus acidophilus | TPY |
| Sulfobacillus benefaciens | AMDSBA4 |
| Sulfobacillus thermosulfidooxidans | Cutipay |
| Variovorax paradoxus | S110 |
| Xanthobacter autotrophicus | Py2 |
